# Identification of Small Molecule Inhibitors of PPM1D Using a Novel Drug Discovery Platform

**DOI:** 10.1101/2024.05.20.595001

**Authors:** Wei Jiang, Subrata Shaw, Jason Rush, Nancy Dumont, John Kim, Ritu Singh, Adam Skepner, Carol Khodier, Cerise Raffier, Ni Yan, Cameron Schluter, Xiao Yu, Mateusz Szuchnicki, Murugappan Sathappa, Josephine Kahn, Adam S. Sperling, David C. McKinney, Alexandra E. Gould, Colin W. Garvie, Peter G. Miller

## Abstract

Protein phosphatase, Mg^2+^/Mn^2+^ dependent 1D (PPM1D), is a serine/threonine phosphatase that is recurrently activated in cancer, regulates the DNA damage response (DDR), and suppresses the activation of p53. Consistent with its oncogenic properties, genetic loss or pharmacologic inhibition of PPM1D impairs tumor growth and sensitizes cancer cells to cytotoxic therapies in a wide range of preclinical models. Given the therapeutic potential of targeting PPM1D specifically and the DDR and p53 pathway more generally, we sought to deepen our biological understanding of PPM1D as a drug target and determine how PPM1D inhibition differs from other therapeutic approaches to activate the DDR. We performed a high throughput screen to identify new allosteric inhibitors of PPM1D, then generated and optimized a suite of enzymatic, cell-based, and *in vivo* pharmacokinetic and pharmacodynamic assays to drive medicinal chemistry efforts and to further interrogate the biology of PPM1D. Importantly, this drug discovery platform can be readily adapted to broadly study the DDR and p53. We identified compounds distinct from previously reported allosteric inhibitors and showed *in vivo* on-target activity. Our data suggest that the biological effects of inhibiting PPM1D are distinct from inhibitors of the MDM2-p53 interaction and standard cytotoxic chemotherapies. These differences also highlight the potential therapeutic contexts in which targeting PPM1D would be most valuable. Therefore, our studies have identified a series of new PPM1D inhibitors, generated a suite of *in vitro* and *in vivo* assays that can be broadly used to interrogate the DDR, and provided important new insights into PPM1D as a drug target.

## INTRODUCTION

Protein phosphatase, Mg^2+^/Mn^2+^ dependent 1D (PPM1D), is a serine/threonine phosphatase that has emerged as a critical regulator of the DNA damage response (DDR) and p53 activation. *PPM1D* is recurrently activated in a wide range of oncologic contexts through amplification, C-terminal truncation mutations, and overexpression.^1,2^ Activation of PPM1D has been associated with various clinical features in cancer, including advanced tumor stage, metastatic potential, therapy resistance, and increased mortality.^3-5^ In multiple preclinical models, PPM1D activation augments malignant transformation and therapy resistance through several mechanisms, including modulation of signaling pathways (p53, p38, NF-KB, mTOR) and changes in cellular processes (DNA repair, cell cycle progression, apoptosis).^6-10^ Accordingly, numerous groups, including our own, have shown that loss of PPM1D impairs tumor growth or sensitizes cancer cells to cytotoxic agents in preclinical models of leukemia, lymphoma, breast cancer, neuroblastoma, Ewing Sarcoma, and glioma.^11-15^ The totality of data supporting PPM1D as a drug target has spawned multiple industrial and academic drug discovery efforts. However, none have yielded a compound with the potential for further progression beyond limited preclinical studies.

Though attractive targets, serine/threonine phosphatases have historically proven difficult to selectively inhibit with small molecules due to their frequent activity as holoenzymes and highly conserved catalytic domains.^16^ Given these challenges in developing competitive inhibitors, *Gilmartin et al*. developed a small molecule allosteric inhibitor of PPM1D, GSK2830371, that was highly selective for PPM1D and impaired the cell growth of a variety of *TP53*-wild-type cancer cell lines.^14^ We subsequently found that high-affinity binding of GSK2830371 to PPM1D occurs at a site on the protein that is unique to PPM1D, explaining its selectivity, and that this binding locks the protein into an inactive conformational state, explaining its mechanism of action.^17^ However, *Gilmartin et al*. could not overcome the pharmacokinetic limitations of GSK2830371 and progress the compound for further clinical development. Additional efforts have identified PPM1D inhibitors, yet none have shown meaningful activity *in vivo*.^18,19^

Given the therapeutic potential of targeting PPM1D specifically and the DDR and p53 pathway more generally, we sought to deepen our biological understanding of PPM1D as a drug target, determine how PPM1D inhibition differs from other therapeutic approaches to activate the DDR, establish a drug discovery platform that could be adapted to broadly study modulators of the DDR and p53, and identify new small molecule inhibitors of PPM1D with the potential for development into clinical agents. We performed small molecule screens, identified hits, and optimized our lead series using medicinal chemistry, pharmacokinetic analyses, and pharmacodynamic studies. In so doing, we developed novel enzymatic, cellular, and *in vivo* assays to study PPM1D and p53 pathway modulators and ultimately identified a series of compounds with comparable *in vivo* activity to GSK2830371 yet distinct chemical properties, allowing for future development efforts. We also deepened our understanding of how PPM1D inhibition activates the DDR differently from other therapeutic approaches and created a drug discovery framework broadly applicable to developing drugs that target the DNA damage response and p53 pathway.

## RESULTS

### Development of a Fluorescent Displacement Assay

Our ultimate goal was to identify compounds that were selective inhibitors of PPM1D. We had previously characterized the allosteric binding activity of GSK2830371 and leveraged this understanding of its biochemical and biophysical properties into a new screening campaign to identify compounds that bound to PPM1D at the same allosteric site as GSK2830371 but with a chemical scaffold that would enable chemical modifications to enhance *in vivo* activity and physicochemical properties.^17^ We, therefore, developed a high-throughput fluorescent polarization assay in which we screened for compounds able to displace an analog of GSK2830371 modified with a fluorescent probe from PPM1D. First, we generated recombinant PPM1D containing the first 420 amino acids (PPM1D_1-420_), as this truncated form is readily produced, stable, and present in a variety of malignancies that harbor C-terminal truncating mutations in *PPM1D* (**Supplemental Figure 1A**).^20^

Next, we generated a fluorescently labeled analog of GSK2830371 that had a moderate binding affinity towards PPM1D_1-420_ and could be displaced by low micromolar binders. We analyzed the properties of the analogs and their fluorophore-modified form towards PPM1D using three assays: the fluorescein diphosphate (FDP) enzymatic assay, a direct binding SPR assay, and the FP binding assay. Briefly, the FDP enzymatic assay measured the rate of cleavage of FDP, an artificial substrate for PPM1D that fluoresces when dephosphorylated and was used to determine the IC_50_ for the analogs (**Supplemental Figure 1B**).^14^ The SPR direct binding assay was used to measure the K_D_ of the analogs for PPM1D_1-420_, which was immobilized on a Ni-NTA chip surface through a dual His(8)-tagged sequence located at the N-terminus. Finally, the FP assay was used as an orthogonal way to determine the K_D_ of binding of the fluorophore-modified form of the analog. After obtaining the desired probe, we developed the FP displacement assay to measure the ability of compounds of varying affinity to displace the analog. Screening analogs of GSK2830371, types of fluorophore, and variation in the linker between the compound and fluorophore yielded Compound 13b-TAMRA (**Figure 1A, Supplemental Figure 1C**), which could inhibit PPM1D in a similar manner as GSK2830371 but could be more readily displaced in the FP assay (**Figure 1B**).

**Figure 1.**
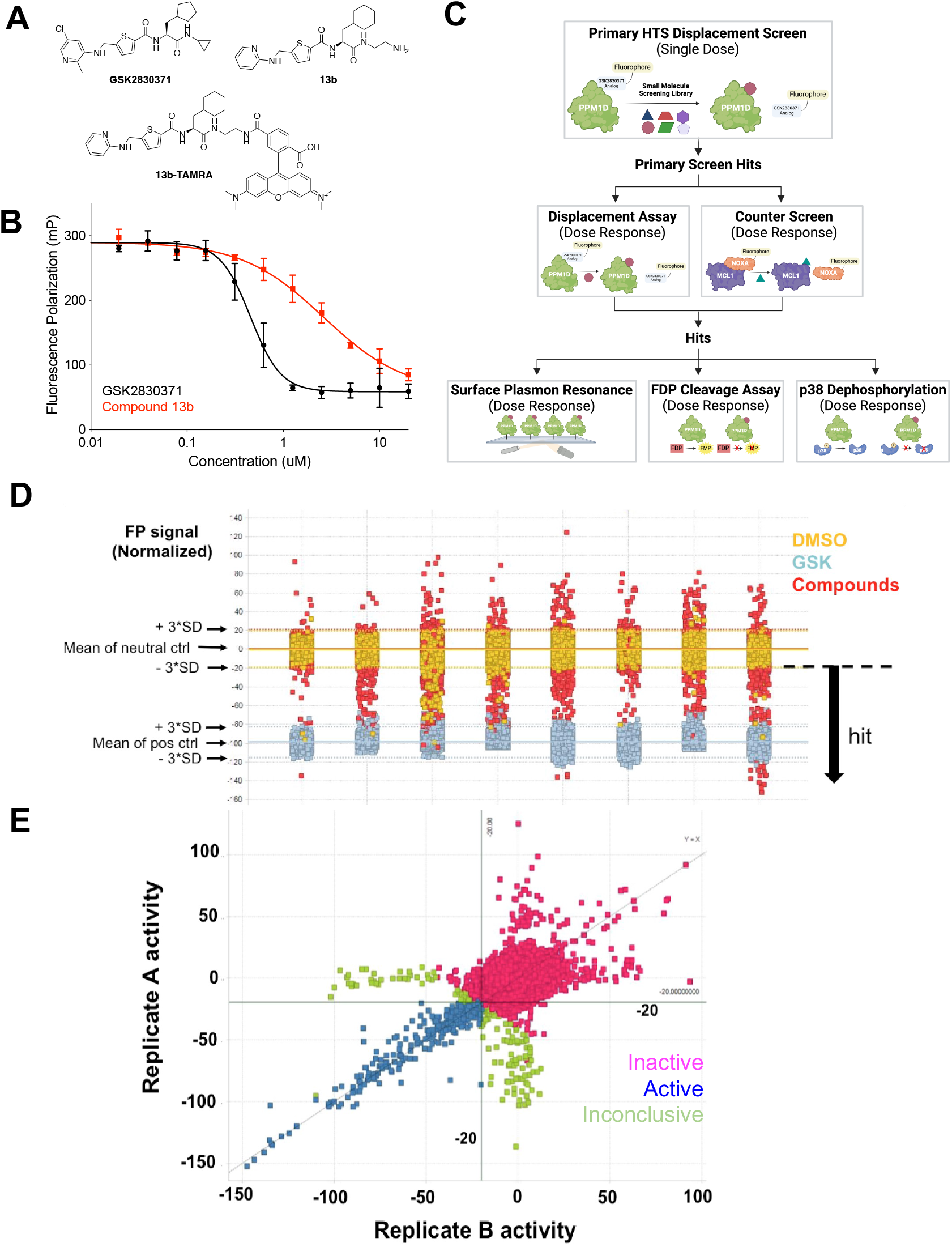
Identification of PPM1D Inhibitors. (A) Structure of GSK2830371, lower affinity analog 13b and fluorescent probe (13b-TAMRA) (B) activity in fluorescent polarization (FP) assay. (C) Schematic of screening cascade. Hits from the primary displacement assay were run in dose-response in the displacement assay and a counter-screen to remove non-specific fluorescent artifacts. These compounds were then screened by surface plasmon resonance, FDP cleavage, and p38 dephosphorylation assays. (D) Summary data from the primary fluorescent polarization screen showing cutoffs for hits, where each column of data represents a set of compounds (the entire screen was run over eight days). (E) Performance of hits across replicates.

### Identification, Validation, and Initial Optimization of PPM1D Inhibitors

We adapted the fluorescence polarization assay to a 1536-well format to screen two chemical libraries (**Figure 1C**). The first was an internal Broad Institute library comprising 200,000 compounds derived from commercial sources and the diversity-oriented synthesis (DOS) library, initially designed to interrogate diverse chemical space.^21^ The second screen was performed at Evotec, Inc. with their Evotec Collection library of 434,963 compounds.^*22*^ The initial round of both screens was performed with a single dose (25 µM) of the compound in duplicates, and we selected compounds that reduced the fluorescence polarization signal by at least three standard deviations compared to the neutral, DMSO control in multiple replicates (**Figures 1D-E**). In parallel, to remove fluorescence artifacts, we excluded compounds that showed an EC_50_ of less than 50 µM in a counter-screen of an unrelated fluorescence polarization assay between MCL1 and a TAMRA-labeled NOXA peptide (data not shown).^23^

We prioritized 256 hits for validation and testing in the SPR and FDP enzymatic assays in addition to a second enzymatic assay that measures the dephosphorylation of p38 MAPK, an endogenous substrate of PPM1D. PPM1D is incubated with full-length p38 MAPK phosphorylated on threonine 180 in this assay. Dephosphorylation of T180 by PPM1D is then measured using liquid chromatography-mass spectroscopy (LC-MS).^17^ The results from these assays ultimately yielded two lead series with distinct scaffolds (thiadiazolo-pyrimidinone **1** and sulfonamide **3**), which we further modified to produce compounds that inhibited PPM1D at single-digit µM concentrations, **2** and **4** (**Figure 2**). By merging features from these two series, we generated racemic cyclohexyl alanine **5**, that exhibited a 100-fold increase in potency (**Supplemental Figure 2A**). We synthesized the individual enantiomers of **5** and were struck to find that the S-enantiomer BRD4761(**6**) was ∼1000-fold more active than the R-enantiomer (**7**), which showed very little activity (**Figure 2**). Initial exploration of the structure-activity relationships (SAR) around BRD4761 led to the replacement of the phenyl sulfonamide amide tail with a di-fluoromethyl phenyl amide, resulting in BRD5049 (**8**) which was then tested across a large panel of human serine/threonine and tyrosine phosphatases, and confirmed to be selective for PPM1D (**Supplemental Figure 2B**).

**Figure 2.**
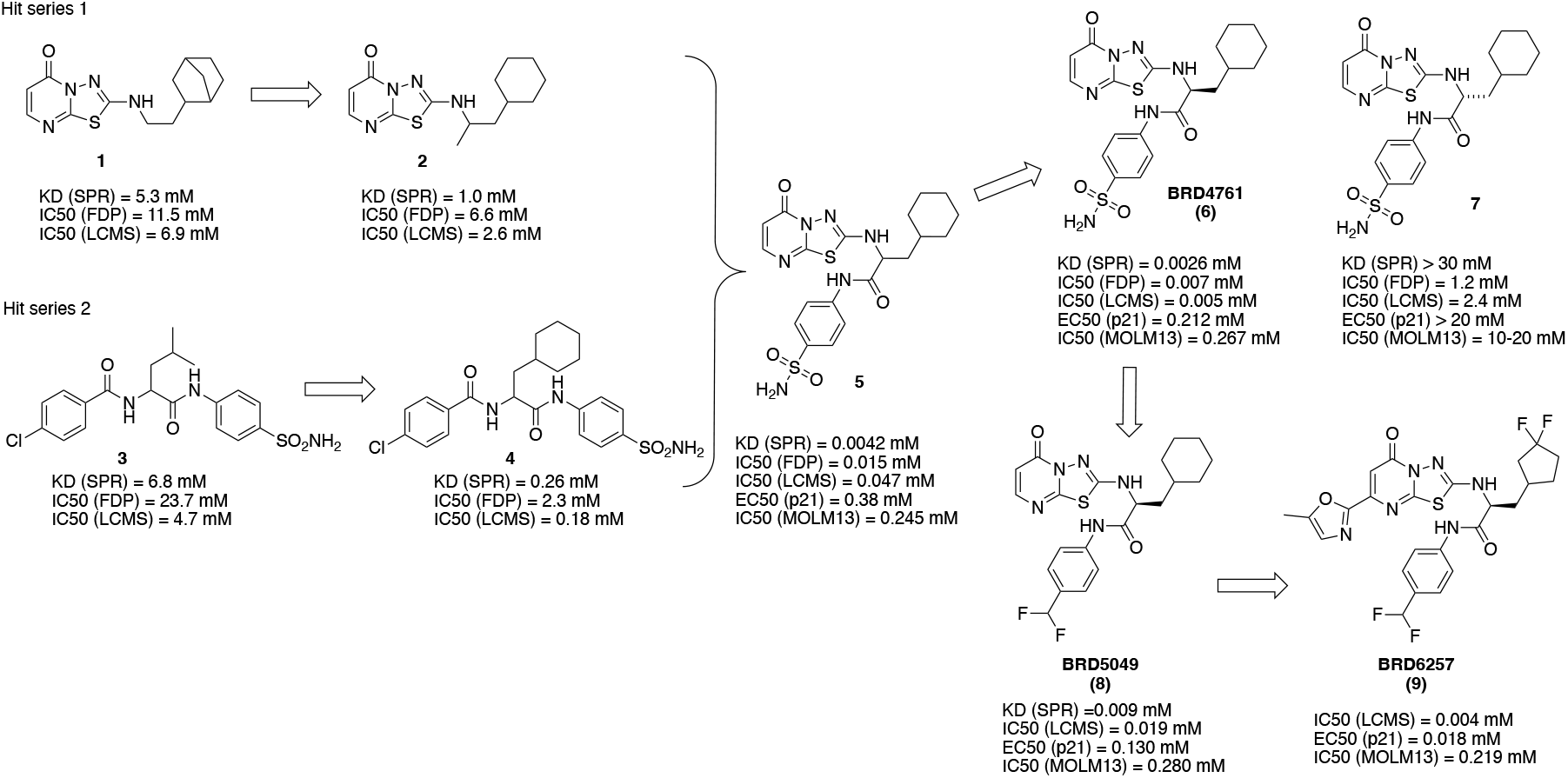
Generation of Lead Compounds. Two hit series were derived from the primary high throughput screening assay and validation studies. These hits were merged to generate **5**, a racemic mixture of active (BRD4761) and inactive 7 isomers. BRD4761 was subsequently modified to create BRD5049 and BRD6257.

### Lead Optimization Using Cell-Based Assays

To further the evaluation of our compounds, we generated cell-based assays to study PPM1D, the DDR, and p53 in distinct ways: dephosphorylation of DDR proteins and PPM1D substrates, activation of p53 and its transcriptional targets, and changes in cell viability (**Figure 3A**). To directly assess the PPM1D-mediated dephosphorylation of DDR proteins, we first established how different small molecule activators of the DDR alter signaling. We treated MOLM13 cells carrying a truncated *PPM1D* (MOLM13-tPPM1D) with DMSO, doxorubicin (DNA damaging agent), AMG232 (an inhibitor of the MDM2-p53 interaction), GSK2830371, or BRD5049 (**Figure 3B**). We observed that doxorubicin strongly induced phosphorylation of DDR pathway proteins, including CHK1, CHK2, and p53, whereas AMG232 only induced p-p53, which is consistent with its mechanism of releasing p53 from MDM2 downstream of CHK1/CHK2. Neither GSK2830371 nor BRD5049 induced phosphorylation of CHK1, whereas both led to a time-dependent increase in p-CHK2, consistent with prior data showing that PPM1D directly dephosphorylates CHK2. In contrast to doxorubicin and AMG232, we did not observe a clear induction of p-p53 with PPM1D inhibition by western blot; however, we did observe an increase in levels of p21 and to a lesser extent BAX and PUMA, all downstream targets of p53, suggesting that PPM1D inhibitors activate p53 signaling in MOLM13 cells, as expected.

**Figure 3.**
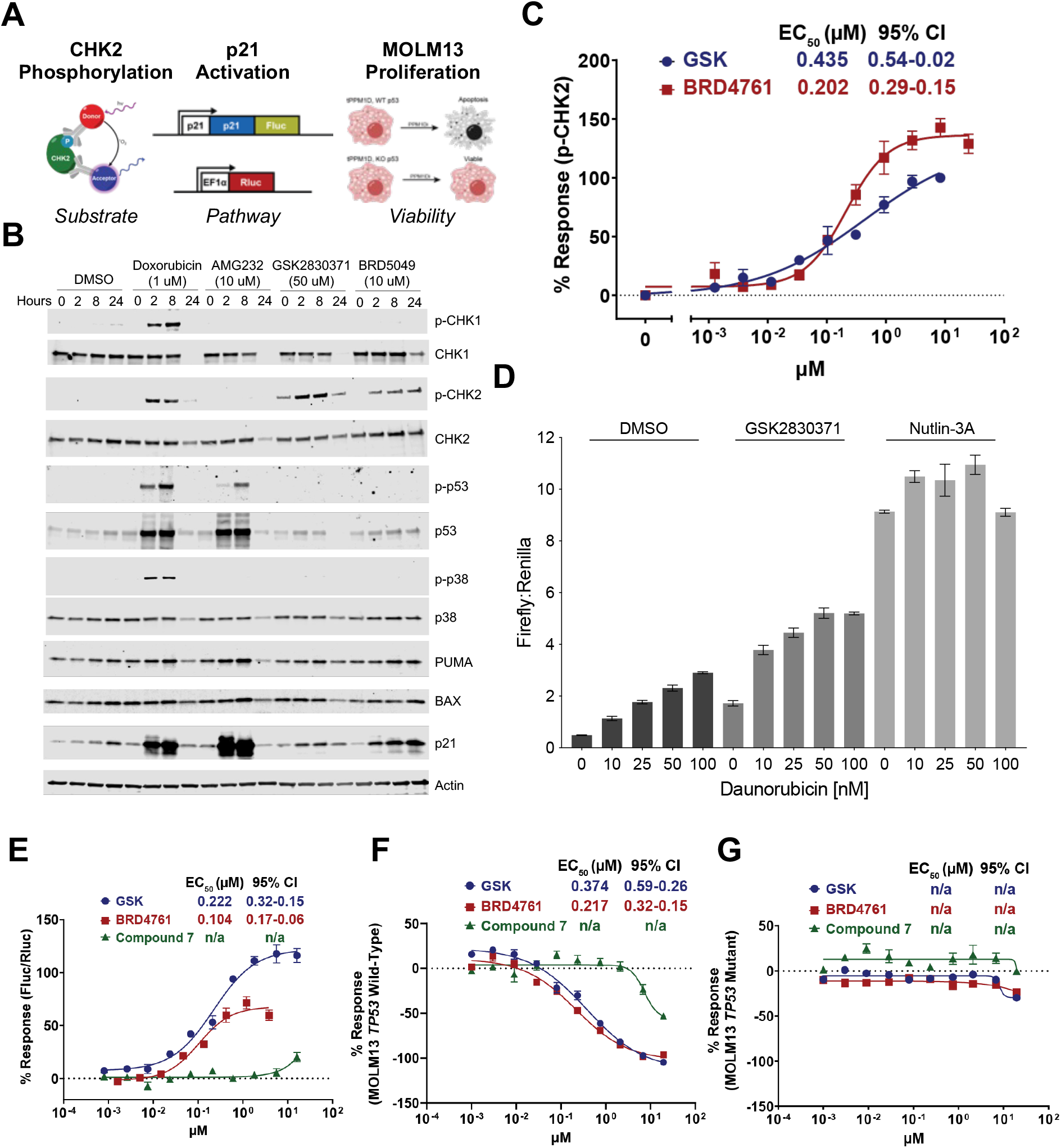
Lead optimization using cell-based assays. (A) Schematic of cell-based assays. (B) Western blot analysis of MOLM13 cells treated with daunorubicin, AMG232, GSK2830371, and BRD5049. (C) Activity of GSK2830371 and BRD4761 in p-CHK2 Alphalisa (D) Firefly:Renilla luciferase level of K562 p21-reporter cells treated with daunorubicin and DMSO, GSK2830371, or Nutlin-3A. (E) Activity of lead compounds in p21-reporter assay. (F-G) Activity of lead compounds in MOLM13 viability assay in *TP53* wildtype (F) and *TP53* mutant (G) cells.

Based on these data, we focused on CHK2 to develop an assay directly measuring PPM1D catalytic activity in cells. To scale our observations, we utilized a commercially available AlphaLISA immunoassay to detect intracellular changes in p-CHK2 in a 384-well format in MOLM13-tPPM1D cells. Consistent with our pilot studies, we found a dose-dependent increase in p-CHK2 upon treatment with GSK283037 and BRD4761. BRD4761 exhibited an EC_50_ similar to GSK2830371 with a slightly greater E_max_ (**Figure 3C**).

Next, to develop an additional readout of p53 pathway activation, we leveraged our previous finding that PPM1D-mediated suppression of p53 decreases CDKN1A (p21) expression, a transcriptional target of p53 (**Supplemental Figure 3**).^17^ Using a K562 cell line with wildtype *TP53*, we utilized a CRISPR/Cas9 homology-directed repair approach to introduce firefly luciferase (ffLuc) into the C-terminus of *CDKN1A*, resulting in the expression of p21-ffLuc upon activation of p53.^24^ For normalization, we introduced a lentiviral vector encoding renilla luciferase (rLuc) driven by an EF1a promoter to express rLuc in these cells constitutively. This system thereby allows for the assessment of p21 levels using the Dual-Glo luciferase assay to quantify the ratio of ffLuc:rLuc, which we confirmed using daunorubicin alone or in combination with either GSK2830371 or Nutlin-3A, an inhibitor of the MDM2-p53 interaction (**Figure 3D**). We adapted this to a 384-well format and confirmed that inhibition of PPM1D by GSK2830371 and BRD4761 both increased ffLuc:rLuc in a dose-dependent fashion, whereas distomer **7** was essentially inactive (**Figure 3E**).

Finally, to measure cellular viability upon PPM1D inhibition, we leveraged an isogenic cell line system. We have previously shown that *TP53* inactivation in MOLM13 cells renders them largely resistant to PPM1D inhibition.^13,25^ For our drug discovery efforts, we utilized CRISPR/Cas9 to generate isogenic cell lines: MOLM13-*tPPM1D TP53* wildtype and MOLM13-*tPPM1D*-*TP53* knockout (KO). Thus, molecules that decrease the viability of MOLM13-*tPPM1D* but not MOLM13-*tPPM1D*-*TP53-KO* are doing so via PPM1D inhibition. In contrast, molecules that decrease the viability of both cell lines presumably do so through a different mechanism, a hypothesis we confirmed with GSK2830371 (**Figures 3F-G**). As expected, both GSK2830371 and BRD4761 exhibited a dose-dependent effect on cell viability of MOLM13-*tPPM1D* but not MOLM13-*tPPM1D*-*TP53-KO* cells, again distomer **7** showed significantly less activity. These data showed that our lead compound, BRD4761, exhibited on-target PPM1D inhibition with efficacy similar to GSK2830371 in cell-based assays despite a significantly different chemical structure.

### Pharmacokinetic Characterization of Lead Compounds

We and others have shown that GSK2830371 has limited *in vivo* activity due to poor pharmacokinetic (PK) characteristics.^14^ We, therefore, prioritized *in vitro* and *in vivo* PK assays to guide further lead optimization. First, we utilized a suite of *in vitro* absorption, distribution, metabolism, and excretion (ADME) assays, including aqueous solubility, plasma stability, clearance in mouse liver microsomes and hepatocytes, permeability and efflux studies in Caco-2 cells, and plasma protein binding (**Table 1**). By generating hundreds of modifications, we developed an understanding of the structure-activity relationship (SAR) of BRD4761 and the drivers of PPM1D potency and ADME properties (**Supplemental Figure 4A**). For example, we found that the bicyclic thiadiazole-pyrimidinone moiety of BRD4761 was important for potency, the phenyl sulfonamide/difluormethyl phenyl substituent could be modified to impact solubility, permeability, and efflux, and alterations to the cyclohexane substituent impacted both potency and metabolic stability (**Supplemental Figure 4B**). We used these data to drive our chemistry efforts, ultimately yielding BRD6257 (**Figure 2**), a compound with relatively better potency in cells and a somewhat improved in vitro ADME profile compared to BRD4761.

**Table 1.**
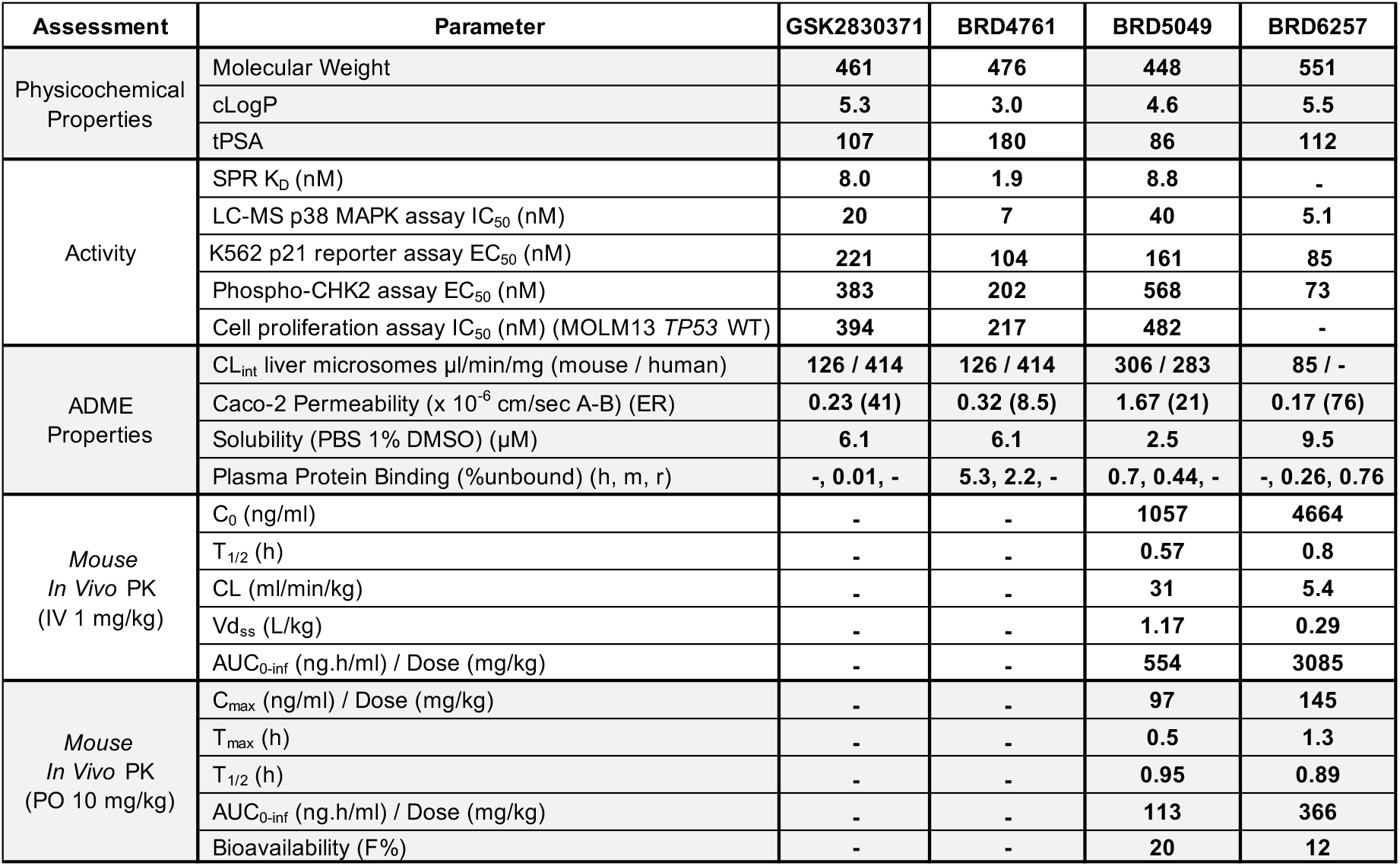
*In Vitro* and *In Vivo* Pharmacokinetic Properties of Lead Compounds.

Next, we performed *in vivo* mouse PK studies to assess compound half-life, clearance rate, and bioavailability with IV and PO formulations of BRD5049 and BRD6257 (**Table 1**). Intravenous administration of a 1 mg/kg dose of BRD5049 and BRD6257 into BALB/c Nude mice resulted in an *in vivo* clearance of 31 and 5.4 ml/min/kg, and area under the curve (AUC) of 554 and 3085 ng.h/ml, respectively. A 5.7-fold decrease in clearance with BRD6257 resulted in a corresponding increase in AUC relative to BRD5049. Oral administration of a 10 mg/kg dose of BRD5049 and BRD6257 resulted in a dose-normalized AUC of 113 and 336 ng.h/ml and bioavailability of 20% and 12%, respectively. After oral administration, plasma exposure with BRD6257 was 3-fold higher than that achieved with BRD5049. Based on these results, we decided to proceed with BRD6257 for *in vivo* pharmacodynamic studies.

### *In Vivo* Pharmacodynamic Evaluation of Lead Compound

Finally, we tested the *in vivo* pharmacodynamic properties of our compounds. To rapidly evaluate the activity of PPM1D inhibitors *in vivo*, we leveraged our K562-p21-ffLuc cell-based reporter assay and adapted it for *in vivo* use, allowing us to monitor on-target pathway modulation in real-time in mice using live bioluminescence imaging (**Figure 4A**). First, we tested the *in vivo* growth characteristics of five single-cell clones derived from the K562-p21-ffLuc parental cell line to avoid experiment-to-experiment variability. We injected one million cells of each clone into both flanks of NRG immunocompromised mice and measured tumor growth.^26^ We chose the clone with the most consistent engraftment and growth (**Supplemental Figures 5A-B**).

**Figure 4.**
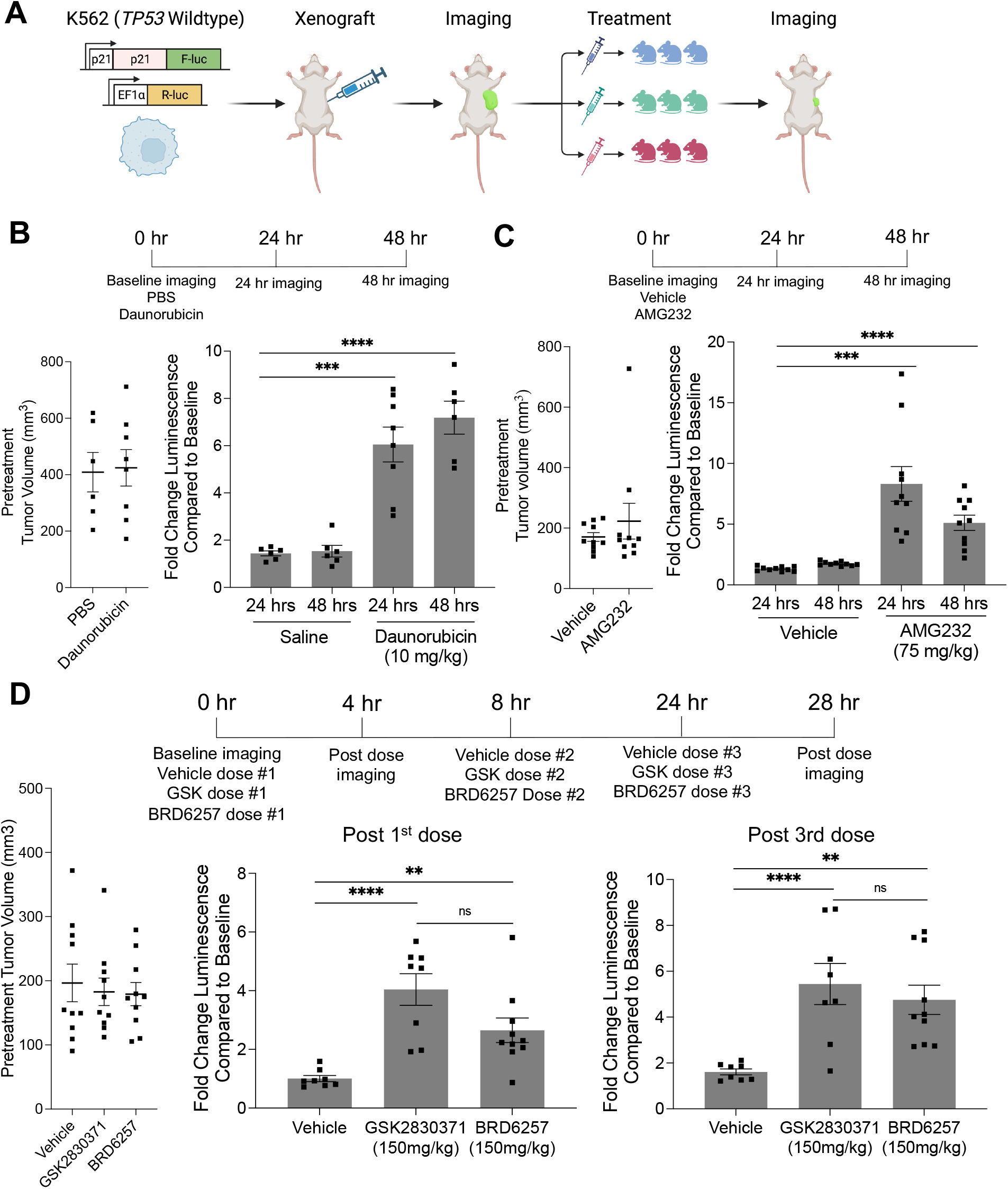
Pharmacokinetic optimization and pharmacodynamic evaluation *in vivo*. (A) Schematic of *in vivo* p21-luciferase assay. Effects of daunorubicin (B) and AMG232 (C) on p21 activation *in vivo* showing pretreatment tumor volume (left) and change in p21-reporter levels (firefly) at given time points (right). (D) Effects of vehicle, GSK2830371, or BRD6257 on p21 activation *in vivo* showing pretreatment tumor volume in each group (left) and change in p21-reporter levels after the 1^st^ and 3^rd^ doses.

Next, to validate the ability of this system to measure p53 activation, we established tumors in mice and confirmed similar tumor sizes between groups on day 13 after inoculation (**Figure 4B**). We then injected the mice with the luciferin reagent, measured p21-luminescent flux from the tumors, and found very high agreement between groups with signal stabilization and stability during the 8-14 minutes after injection (**Supplemental Figure 5C**). Next, we dosed mice with vehicle (phosphate buffered saline) or daunorubicin (10mg/kg) by intraperitoneal injection and observed increases in p21-luciferase levels consistent with p53 pathway activation at 24 and 48 hours after dosing (**Figure 4B and Supplemental Figures 5C-D**). We also used quantitative PCR to quantify the expression of *CDKN1A* (p21) in explanted tumors at 48 hours and found a small but significant increase in the daunorubicin-treated group. However, the effect was less than the difference observed using the p21-luciferase-based readout (**Supplemental Figure 5E**). To test the system with an orthogonal mode of p53 activation, we utilized AMG232, which is orally bioavailable.^27^ Using a similar experimental design, we confirmed equal tumor size and bioluminescent levels at baseline, then dosed animals with either vehicle (5% DMSO, 20% Kolliphor EL, 75% water) or AMG232 (75 mg/kg) (**Figure 4C and Supplemental Figure 5F**). We imaged the animals after 24 and 48 hours and observed a significant induction in p21-luciferase levels in the AMG232-dosed mice compared to vehicle control, consistent with activation of the p53 pathway (**Figure 4C**).

Having established a system for rapid interrogation of the p53 pathway *in vivo*, we next sought to test the effects of PPM1D inhibition. We established tumors and separated the animals into three groups: vehicle (5% DMSO, 20% Cremophor EL, 75% water), GSK2830371 (150mg/kg), and BRD6257 (150mg/kg). We confirmed that the baseline body weights, tumor volumes, and luminescence levels were similar across groups (**Supplemental Figure 5G-H**). Next, the animals were dosed three times by oral gavage (0 hours, 8 hours, and 24 hours) and imaged four hours after the first and last doses. Relative to baseline luminescence, we observed a significant increase in p21-luciferase with 150 mg/kg GSK2830371 and 150 mg/kg BRD6257 relative to the vehicle. Four hours after a single dose, we observed a 4-fold (GSK2830371) and 2.7-fold (BRD6257) increase over the vehicle, and four hours after the last dose, we observed a 5.4-fold (GSK2830371), 4.8-fold (BRD6257), and 1.6-fold change over baseline vehicle (**Figure 4D**). Though the average induction in the GSK2830371 group was slightly higher than in the BRD6257 group, the difference was not statistically significant. As expected from the mechanism of action and the results of our biochemical studies, the increase in p21 levels was higher in the daunorubicin and AMG232 groups than in the PPM1D inhibitor groups. These data confirm that oral administration of BRD6257 inhibits PPM1D and activates the p53 pathway *in vivo* comparable to GSK2830371.

## DISCUSSION

Modulating the DNA damage response has emerged as an area of great therapeutic interest. Within this framework, we have focused on *PPM1D*, a gene recurrently activated across multiple oncologic contexts with extensive preclinical validation as a drug target in cancer. Our ultimate goal was to identify new small-molecule inhibitors of PPM1D. In so doing, we sought to deepen our biological understanding of PPM1D as a drug target and how it compares to other approaches to activate the DDR and p53. To address these questions, we established and deployed a suite of enzymatic, biochemical, cellular, and *in vivo* assays. Importantly, these systems can be readily adapted to study the DDR and facilitate drug discovery efforts beyond PPM1D.

Our drug discovery efforts focused on developing inhibitors with four primary characteristics: (1) chemical distinction from published compounds, (2) selectivity for PPM1D, (3) activity *in vivo*, and (4) improved PK over GSK2830371. To this end, we screened for compounds that displaced a GSK2830371 analog from its allosteric binding site on PPM1D. Using established and new enzymatic, biochemical, and cell-based assays, we optimized and tested our lead compounds in pharmacokinetic and pharmacodynamic assays. Our most advanced compound, BRD6257, is PPM1D-selective and chemically distinct from published inhibitors (**Figure 5**), yet it displayed *in vivo* activity similar to GSK2830371 following oral administration in mice.

**Figure 5.**
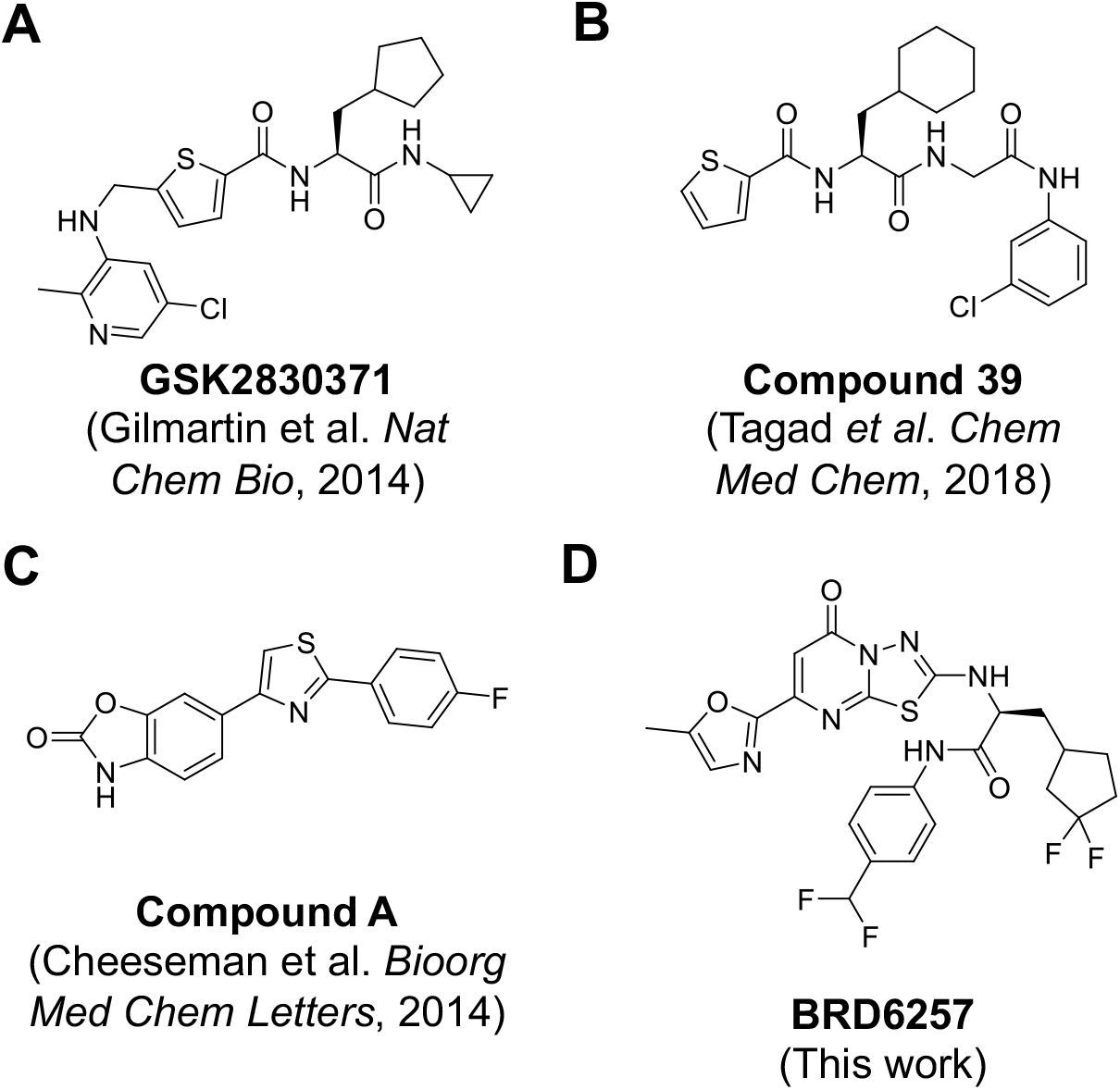
Previously reported PPM1D Inhibitors. PPM1D inhibitors reported from prior screens, none of which have progressed past preclinical studies.

We developed an understanding of the structure-activity relationships of our lead compounds, their activity against PPM1D, and their performance in *in vitro* pharmacokinetic studies. However, despite this knowledge, our modifications that improved pharmacokinetics came at the cost of potency, and we ultimately could not generate a compound with a significantly longer half-life than GSK2830371 while maintaining potency. One potential explanation is that our screening approach only identified compounds that bind to the same allosteric site as GSK2830371, and there may be inherent limitations in PK-efficacy trade-offs of drugs that can do so. Alternative screening approaches to identify compounds that bind to other sites may overcome the challenges posed by our lead series. While we screened over 600,000 compounds that included a commercially available set and a diversity-oriented synthesis library, expanded screening of more extensive compound collections may yield new compounds with distinct chemical and PK profiles. Additionally, despite Gilmartin et al.’s extensive crystallography efforts and our own group’s crystallography and cryo-electron microscopy work, the structure of PPM1D remains unsolved.^17^ Moreover, neither our computational model of PPM1D nor those generated by the most recent iterations of Alphafold yielded the required detail to model the binding of PPM1D to GSK2830371 or our lead compounds. As a result, we were not structurally enabled, which made our medicinal chemistry efforts even more challenging. Therefore, future efforts to solve the structure of PPM1D may provide significant new opportunities for further development of our lead compounds. Finally, while our medicinal chemistry efforts were substantial, we did not explore all potential modifications, and there may be opportunities for further development of our lead series.

Identifying novel PPM1D inhibitors required the development of multiple new assays to assess substrate dephosphorylation, pathway activation, and cell viability. To directly assess PPM1D’s phosphatase activity on known substrates, we developed an enzymatic, mass-spectrometry-based readout of full-length phospho-p38 and utilized a cell-based assay of phospho-CHK2. To measure the effects of PPM1D inhibitors on the p53 pathway, we developed a new, luciferase-based readout of p53 transcriptional activity by genetically modifying the endogenous *CDKN1A* locus. Finally, to test the on and off-target cytotoxic activity of PPM1D inhibitors, we generated and utilized *TP53* isogenic MOLM13 cell lines. While we used these assays to develop PPM1D inhibitors, they can also be readily applied with other modalities (e.g., genetic perturbations) and used to study other members of the DDR and p53.

Traditional PK/PD studies most frequently rely on *ex vivo* analysis of tumors using standard methods such as western blot, quantitative PCR, or immunohistochemistry to quantify changes in downstream targets/pathways. Indeed, we assessed the pharmacodynamic activity of GSK2830371 by western blot analysis of phosphorylation changes in PPM1D targets in explanted tumors.^14^ However, none of these approaches allow for a real-time, *in vivo* quantification of pathway activation. To overcome this problem, we established a xenograft model with our p21-luciferase reporter cell line, which allowed us to measure the induction of p21 in real-time *in vivo* by bioluminescent imaging. We quantified the activation of the DDR and p53 after treatment with the DNA-damaging chemotherapy, daunorubicin, then used this model system to test the effects of GSK2830371 and our lead compound. While we focused on PPM1D modulation, this system can be applied to study and quantitatively compare any agents or perturbations expected to activate the DDR or p53 signaling.

Our data suggest that PPM1D inhibitors are biologically and therapeutically distinct from inhibitors of the MDM2-p53 interaction. PPM1D inhibition activates p53 and induces cell death via enhanced phosphorylation of multiple DDR substrates, including p53. However, consistent with its role as a negative regulator of the DDR, PPM1D may be most therapeutically important in the context of concurrent genotoxic stress. Indeed, we and others have previously shown that PPM1D inhibition can sensitize a series of cancer cells to DNA-damaging chemotherapies and radiation.^11-15^ In contrast, inhibitors of the MDM2-p53 interaction, such as AMG232, prevent MDM2-mediated sequestration and degradation of p53, thereby increasing intracellular levels of activated p53 without concurrent activation of upstream members of the DDR. This difference may explain why AMG232 exhibits a greater cytotoxicity than PPM1D inhibitor monotherapy. While this study did not address toxicity, we have previously shown that genetic loss of *Ppm1d* in adulthood was well tolerated in mice without evidence of impaired platelet production, which contrasts with the thrombocytopenia observed with MDM2-p53 inhibitors.^13,28^ In summary, our studies have identified a series of new PPM1D inhibitors, generated a suite of *in vitro* and *in vivo* assays that can be broadly used to interrogate the DDR and provided important new insights into PPM1D as a drug target.

## ACKNOWLEDGMENTS

This work was supported by grants from the National Institutes of Health (NIH) (K08-CA263181 [P.G.M.]; K08-CA252174 [A.S.S.]), the Edward P. Evans Foundation (P.G.M.), and the Department of Defense (DOD) (CA220652 [P.G.M.]; CA210827 [A.S.S.]). The authors would like to thank Deerfield Ventures for supporting the work. The authors thank Dr. Benjamin Ebert for his thoughtful advice and feedback throughout this study.

## AUTHOR CONTRIBUTIONS

P.G.M. and C.G. designed and conceived the study; W.J, S.S., J.R., N.D., J.K., R.S., A.S., C.K., C.R., X.Y., M.S., N.Y., C.S., and M.S. performed the experiments. P.G.M, C.G., W.J, S.S., J.R., A.S.S., J.K., N.D., R.S., S.G. analyzed and interpreted the data. P.G.M. and C.G. drafted the manuscript.

## METHODS

### Protein production for screening and characterization

The production of PPM1D(1-420), p38 MAPK with the stabilizing mutation C162S, and a constitutively active form of MKK6(S207E, T211E) have been described in detail previously.^29,30^ In brief, the genes for all three proteins were codon-optimized and were expressed and purified from *E. coli*. PPM1D(1-420) was expressed with an N-terminal tag comprised of a polyhistidine sequence followed by the SUMO protein. p38 and MKK6 were expressed with an N-terminal tag consisting of a polyhistidine followed by a TEV cleavage site. The proteins were purified by a combination of metal chelating affinity chromatography and size exclusion chromatography, with the tag removed during purification. For the SPR assay, PPM1D(1-420) was expressed with an N-terminal His(8)-28aa-His(8) affinity tag in order to immobilize on an Ni-NTA chip. Expression and purification were performed in a similar manner as described above, except the N-terminal tag was not removed. The phosphorylated form of p38 MAPK was generated by incubating with MKK6 in the presence of ATP.

### Fluorescence Polarization (FP) Displacement Assay and Counterscreen

The FP assay was performed in buffer containing 50 mM TRIS pH 7.5, 50 mM NaCl, 1 mM MgCl2, 0.01% P20, 2% DMSO. The assay was run in 1536-well plates (Aurora #ABI040100A) in a final volume of 8 uL per well. The reaction was initiated by the addition of a solution containing 400 nM PPM1D and a 5 nM fluorescence probe to the plate, where compounds at various concentrations were pre-dispensed. The plate was then incubated at room temperature for 1 hour with the lid on, and subjected to data collection by EnVision Multimode Plate Reader at Ex/Em 544/579 nm. The FP counterscreen was carried out, as previously described, using 1 uM MCL1 and 25 nM Noxa peptide-TAMRA as the probe in the buffer containing 25 mM Hepes, 100 mM NaCl, 0.005% P20, pH 7.4.^23^

### Surface Plasmon Resonance (SPR) assay

The SPR assay was performed on a Biacore 200 using Series S sensor chip NTA following the reported protocol.^17^ Double-His_8_-tagged PPM1D_1-420_ and/or PPM1K_89-351_ were immobilized. All compounds were first analyzed using the classic multi-cycle setup. For those compounds showing slow kinetics and strong potency in the enzymatic assays, a second round of SPR using a single-cycle setup was performed. The chip surface needed to be re-generated after each compound injection under a single-cycle setup. K_D_, *k*_on,_ and *k*_off_ were then calculated.

### Fluorescein Diphosphate (FDP) Enzyme Activity Assay

The enzymatic assay of PPM1D, PPM1A, or PPM1K using the artificial substrate, fluorescein diphosphate (FDP), was carried out following the reported protocol.^14^ Cleavage of FDP by the enzyme yields fluorescein monophosphate and fluorescein, resulting in an increase in fluorescence signal. The enzyme was first incubated with compounds at various concentrations for 30 min, followed by the addition of FDP and kinetic fluorescence intensity (FI, 485/530 nm) data collection for 15 min. The extracted slope indicates the enzyme activity.

### p38 MAPK Enzyme Activity Assay

We developed an MS-based enzymatic assay to measure the dephosphorylation of the PPM1D substrate p38 MAPK.^17^ In this assay, PPM1D was first incubated with compounds for 30 min, followed by the addition of diphosphorylated p38 MAPK and incubation at R.T. for two h. The reaction was quenched, and BSA was added as a carrier protein and injection control for LC-MS. Data collection was executed on the same day using multiple-reaction monitoring (MRM) using a UPLC/MS system.

### Phosphatase Selectivity Assay

The phosphatase selectivity assay was performed on the Eurofins Discovery Phosphatase Profiler (https://www.eurofinsdiscovery.com/solution/phosphatases).

### Cell-Based Assays

#### Western Blot Analysis

MOLM13 cells were adjusted to 4 million cells/mL in complete media (RPMI + 10% FBS + 1X P/S) and 0.5 mL was distrubted into 6-well plates. Compounds were added as 0.5 mL at 2X the final concentration in complete media. At designated time points, cells were collected, washed with PBS and stored at -80 °C. Cell pellets were thawed and lysed in 75 µL RIPA buffer (25 mM Tris•HCl pH 7.6, 150 mM NaCl, 1% NP-40, 1% sodium deoxycholate, 0.1% SDS, Thermo 89900), 1X benzonase, 1X cOmplete^™^ Protease Inhibitor Cocktail, 1X phosphatase inhibitors (cell signaling #5870). After 10 min on ice, insoluble material was removed by centrifugation at max speed for 10 min at 4 °C. 65 µL of clarified lysate was mixed with 10 µL 0.5 M TCEP and 25 µL 4X loading buffer. A BCA assay was performed on remaining lysate and samples were normalized with 1X SDS loading buffer. 10-30 µg of protein was resolved on Criterion TGX Precast Midi Protein Gels (Bio-Rad) then a wet transfer onto Immobilon-FL (Millipore). Intercept (TBS) blocking buffer (Licor) was used as indicated by the manufacturer for blocking. Antibody identities and dilutions are listed below.

| Epitope | clone | vendor | cat# | species | Dilution | Secondary | Dilution |
| --- | --- | --- | --- | --- | --- | --- | --- |
| p53 | DO-1 | CST | 18032 | mouse mAb | 1:1000 | IRDye® 680RD Goat anti-Mouse | 1:10,000 |
| phos-p53 (S15) |  | CST | 9284 | rabbit pAb | 1:1000 | IRDye® 800CW Goat anti-Rabbit | 1:10,000 |
| p38 |  | CST | 9212 | rabbit pAb | 1:1000 | IRDye® 680RD Goat anti-Rabbit | 1:10,000 |
| phos-p38 (Thr180/Tyr182) | 28B10 | CST | 9216 | mouse mAb | 1:1000 | IRDye® 800CW Goat anti-Mouse | 1:10,000 |
| Chk2 | 1C12 | CST | 3440 | mouse mAb | 1:1000 | IRDye® 680RD Goat anti-Mouse | 1:10,000 |
| phos-Chk2 (Thr68) | C13C1 | CST | 2197 | rabbit mAb | 1:1000 | IRDye® 800CW Goat anti-Rabbit | 1:10,000 |
| Chk1 | G-4 | SantaCruz | sc-8408 | mouse mAb | 1:1000 | IRDye® 680RD Goat anti-Mouse | 1:10,000 |
| phos-Chk1 (Ser345) | 133D3 | CST | 2348 | Rabbit mAb | 1:1000 | IRDye® 800CW Goat anti-Rabbit | 1:10,000 |
| PUMA | E2P7G | CST | 98672 | rabbit mAb | 1:1000 | IRDye® 800CW Goat anti-Rabbit | 1:10,000 |
| BAX | D2E11 | CST | 5023 | rabbit mAb | 1:1000 | IRDye® 800CW Goat anti-Rabbit | 1:10,000 |
| p21 | 12D1 | CST | 2947 | rabbit mAb | 1:1000 | IRDye® 800CW Goat anti-Rabbit | 1:10,000 |
| PPM1D | F-10 | SantaCruz | sc-376257 | mouse mAb | 1:1000 | IRDye® 800CW Goat anti-Mouse | 1:10,000 |
| Actin | AC15 | sigma | A1978 | mouse mAb | 1:20,000 | IRDye® 680RD Goat anti-Mouse | 1:20,000 |

#### p-CHK2 Assay

The Alphalisa p-CHK2 (Thr68) assay (PerkinElmer) was adapted to 384-well format as follows, 60,000 MOLM13-tPPM1D^25^ cells in 20 µL complete media (RPMI + 10% FBS + 1X P/S) were added to the wells of a 384-well plate and incubated at 37 °C for 2 h. A pinning tool was used to transfer compounds to the assay plates and incubation was continued for 4 h. Measurement of p-CHK2 was performed using the AlphaLISA Surefire Ultra Human Phospho-CHK2 (Thr68) Detection Kit (Revity). Briefly, 5 µL of 5X lysis was added to each well and the plates were shaken at RT for 10 min. A mix of reaction buffer 1, reaction buffer 2, activation buffer and acceptor beads were prepared according to the manufacturer and 5 µL was added to each well. The plate was sealed and incubated at RT for 2 h. Dilution buffer and acceptor beads were prepared according to the manufacturer and 5 µL were added to each well. The plate was sealed and incubated at RT for 24 h before the AlphaLISA signal was measured using an Envision platereader (PerkinElmer).

#### p21 Reporter Assay

The p21 luciferase reporter cell line was generated from K562 cells engineered to carry wildtype *TP53*.^24^ To generate the reporter, firefly luciferase was introduced into the C-terminus of the endogenous CDKN1A locus using the endogenous tagging methodology previously described.^31^ Lentivirus encoding renilla luciferase in the pLX307 vector was introduced into these cells, yielding the final reporter cell line. 12,500 cells in 30 µL complete media (RPMI + 10% FBS + 1X P/S) added to 384-well plates containing predispensed compounds (Echo). Afer 24 h levels of firefly luciferase and renilla luciferase in each well were measured using the DualGlo Luciferase Assay System (Promega), and the ratio of firefly luciferase to renilla luciferase was calculated.

#### MOLM13 Viability Assay

The MOLM13 cells isogenic for *TP53* knockout were generated as previously described.^25^ 2,000 MOLM13-tPPM1D cells or MOLM13 cells isogenic for *TP53* knockout cells in 50 µL complete media (RPMI + 10% FBS + 1X P/S) added to 384-well plates containing predispensed compounds (Echo). The plates were incubated for 96 h then equilibrated to RT. 10 µL of CellTiter-Glo reagent (Promega) were added to each well. After incubation with gentle agitation for 10 min at RT, luminescence was measured using an Envision platereader (PerkinElmer). Cells were grown in RPMI + 10% FBS and plated in 384 well format at a density of 2000. After the addition of compounds, the cells were grown for three days, and the viability of each well was assessed using the CellTiter-Glo reagent (Promega).

#### Kinetic Solubility

For the kinetic solubility assessment **t**he compounds were dissolved in DMSO to prepare a 10 mM stock concentration. 495 µL of phosphate buffered saline (PBS, pH 7.4) or DMSO was added to appropriate wells of a 96-well plate followed by 5 µL of 10 mM DMSO stocks of the test compounds. The final concentration of the test compound in DMSO was 100 µM. Samples were prepared in duplicate. The sample plate was sealed and vortexed at 750 rpm for 18 hours on a vortex shaker at room temperature. At the end of the incubation period, the sample plate was centrifuged at 4000 rpm for 15 mins. 200 µL of the supernatant from each well was transferred to a fresh plate followed by addition of the same volume of acetonitrile. Samples were analyzed by HPLC-UV analysis. The peak area of the compound in DMSO was used as a reference to determine the concentration of the compound in PBS.

### Metabolic Stability

The study was conducted to assess the metabolic stability of compounds in mouse and human liver microsomes. A 445 μL aliquot of liver microsomal proteins (0.56 mg/mL) and 5 μL of compound (100 μM) were added to a 96-well microplate and pre-incubated for 5 min at 37 °C. 50 μL of NADPH (10 mM) was added to initiate the reaction. The final concentration of microsomal proteins, compound and NADPH in the incubation was 0.5 mg/ml, 1 μM and 1 mM, respectively. A 60 μL aliquot was transferred to a fresh plate at pre-determined time points 0, 5, 15, 30, 45, and 60 min. The reaction was terminated with 180 μL of quenching solvent containing internal standard. NADPH-free control incubations were carried out along with the main experiment, and the samples were collected at 0 and 60 min. The plates were placed on a shaker for 10 minutes, followed by centrifugation at 4000 rpm for 20 minutes at 4°C. The samples were diluted 3-fold with water, mixed well and analyzed using LC-MS/MS. The percentage of compound remaining at each time point, half-life, intrinsic clearance, and clearance *in vivo* (well-stirred model) was calculated.

### Caco-2 Permeability

Compounds were screened for permeability across human colorectal adenocarcinoma cells (Caco-2 cells) using Transwell plates. Caco-2 cells purchased from ATCC were seeded onto polyethylene membranes (PET) in 96-well Corning Insert plates at a cell density of 1 x 10^5^ cells/ cm^2^ and media refreshed every 4∼5 days until Day 21 for confluent cell monolayer formation. Before the experiment, TEER was measured to check the monolayer integrity. Monolayers with a TEER value above 400 Ω·cm2 were used for the permeability studies. Post experiment, monolayer integrity was tested using Lucifer yellow (LY-100 μg/mL) solution, and percentage LY transfer across the monolayer was detected using a fluorescence method. The apparent permeability and efflux ratio of compounds across Caco2 cell monolayer were determined by performing apical to basolateral (A to B) and basolateral to apical (B to A) permeability. In brief, 5 μM of each compound was spiked in the donor compartment and maintained at 37 °C at 100 rpm on an orbital shaker. Samples were collected from the receiver compartment at 120 min. Samples were collected from the donor compartment at 120 min to assess mass balance. The samples were analyzed using LC-MS/MS.

### Plasma Protein Binding

Rapid equilibrium dialysis (RED) assay was performed using inserts (Thermo Fischer Scientific, product No. 89810) and base plate-made of PTFE material (Thermo Fischer Scientific, product No. 89811). The rapid equilibrium dialysis base plate with inserts was assembled following manufacturer’s instruction. 995 μL of plasma was spiked with 5 μL of the compound (200 μM). 350 μL of buffer was added to the buffer chamber followed by addition of 250 μL of spiked plasma into the sample chamber of the RED insert. The plate was rotated at approximately 450 rpm on an orbital shaker in a humidified incubator with 5% CO2 at 37°C for 4 hours. At the end of the dialysis, aliquots of 50 μL samples from the buffer side and matrix side of the dialysis device were transferred to a fresh 96-well plate. An equal volume of opposite blank matrix (buffer or matrix) in each sample was added to reach a final volume of 100 μL. All samples were further processed by protein precipitation for LC-MS/MS analysis. The fraction unbound was determined as a ratio of the compound peak area in the buffer chamber and the plasma chamber.

### *In Vivo* Pharmacokinetic Studies

Female BALB/c Nude Mice were used for pharmacokinetic assessment. The mice were randomly assigned to two treatment groups (n=3 per group). The compound was dissolved in 5%DMSO/20%Cremophor EL/75%water at the appropriate concentration for dosing. On dosing day, mice received a single intravenous (i.v.) dose (1 mg/kg) or oral (p.o.) dose (10 mg/kg) of test compound, and ∼100 μL blood samples were collected at 0.083 (only for i.v.), 0.25, 0.5, 1, 2, 4, 8, and 24 h after dosing. Plasma was prepared following centrifugation of the collected blood samples. 4 μL of each sample was quenched with 120 µL of acetonitrile containing internal standard followed by vortex-mixing for 10 min at 800 rpm and centrifugation for 15 min at 3220 × g, 4 °C. A fit-for purpose bioanalysis method was developed, and plasma concentration−time profile was obtained using LC-MS/MS. The plasma concentration−time data were subjected to non-compartmental analysis using Phoenix WinNonlin (Version: 6.3) to assess the PK parameters.

### *In Vivo* Assessment of Compound Activity

One million of the K562 luciferase reporter cells were injected bilaterally into the flanks of NRG mice.^27^ 11-13 days later, upon palpable tumor formation, the levels of p21-luciferase activity were quantified by bioluminescent imaging. Tumor sizes were assessed using digital calipers. The mice were anesthetized, injected with 150mg/kg luciferin intraperitoneally, and after 4 minutes the mice were serially imaged on the IVIS SpectrumCT platform. Bioluminescence was quantified using the Living Image software.To assess the effects of treatments in this model, the animals were treated with phosphate buffered saline (intraperitoneal injection), daunorubicin (10mg/kg, intraperitoneal injection), vehicle (5% DMSO, 20% Cremophor EL, 75% H2O, oral gavage), AMG232 (75mg/kg, oral gavage), GSK2830371 (150mg/kg, oral gavage), or BRD6257 (150 mg/kg, oral gavage). The frequency of doses and timepoints of imaging are outlined in **Figure 4**.

