## Supplementary Figures for "Identification of Small Molecule Inhibitors of PPM1D Using a Novel Drug Discovery Platform"

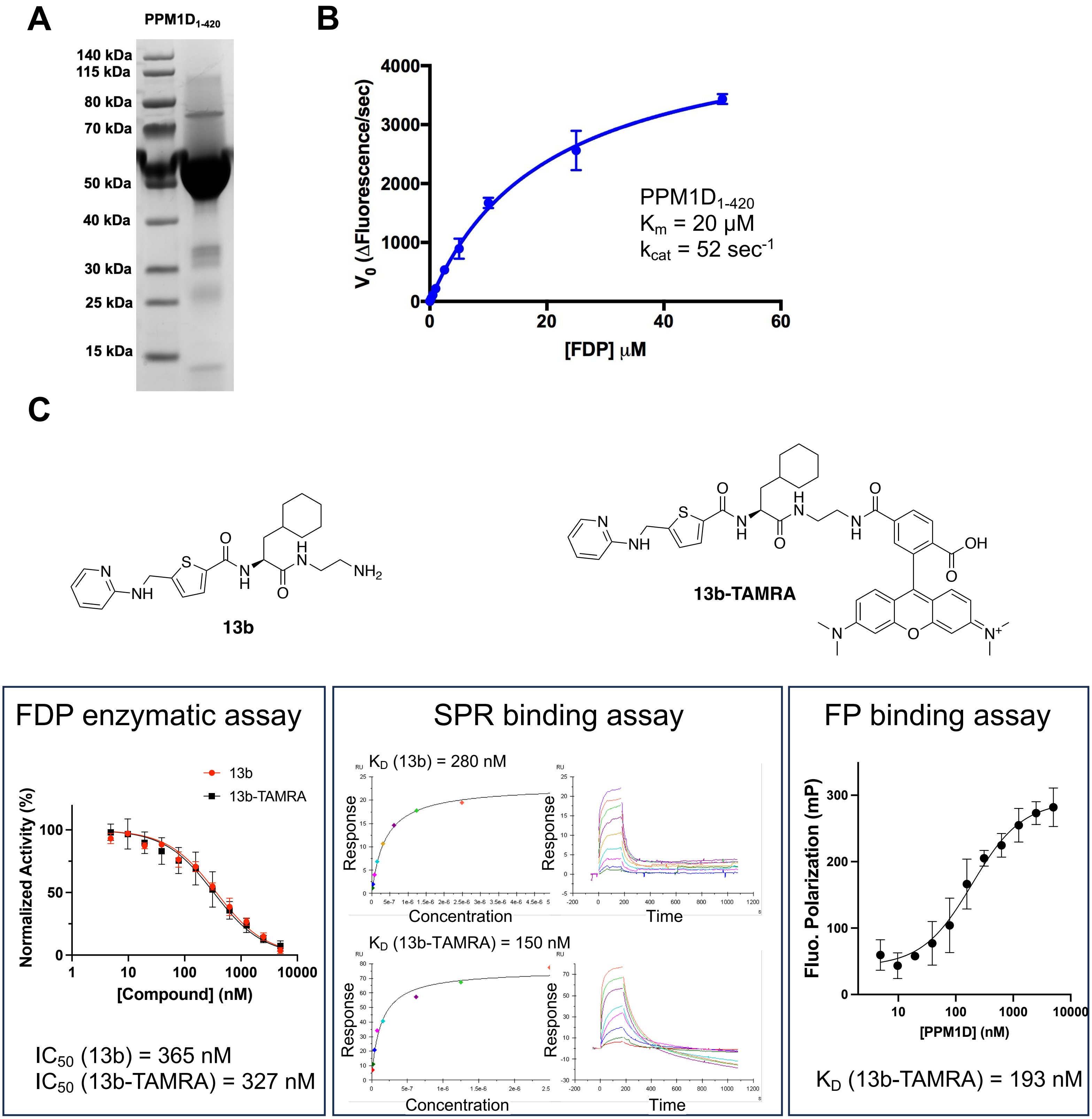


**Supplemental Figure 1. Identification of PPM1D Inhibitors.** (A) SDS-PAGE gel showing generation of recombinant $PPM1D_{1\text{-}420}$ used for subsequent assays. (B) Enzymatic activity of recombinant $PPM1D_{1\text{-}420}$ in FDP assay. (C) Characterization of 13b and 13b-TAMRA using FDP enzymatic assay, SPR direct binding assay, and FP binding assay (applicable to 13-TAMRA only).

**A**

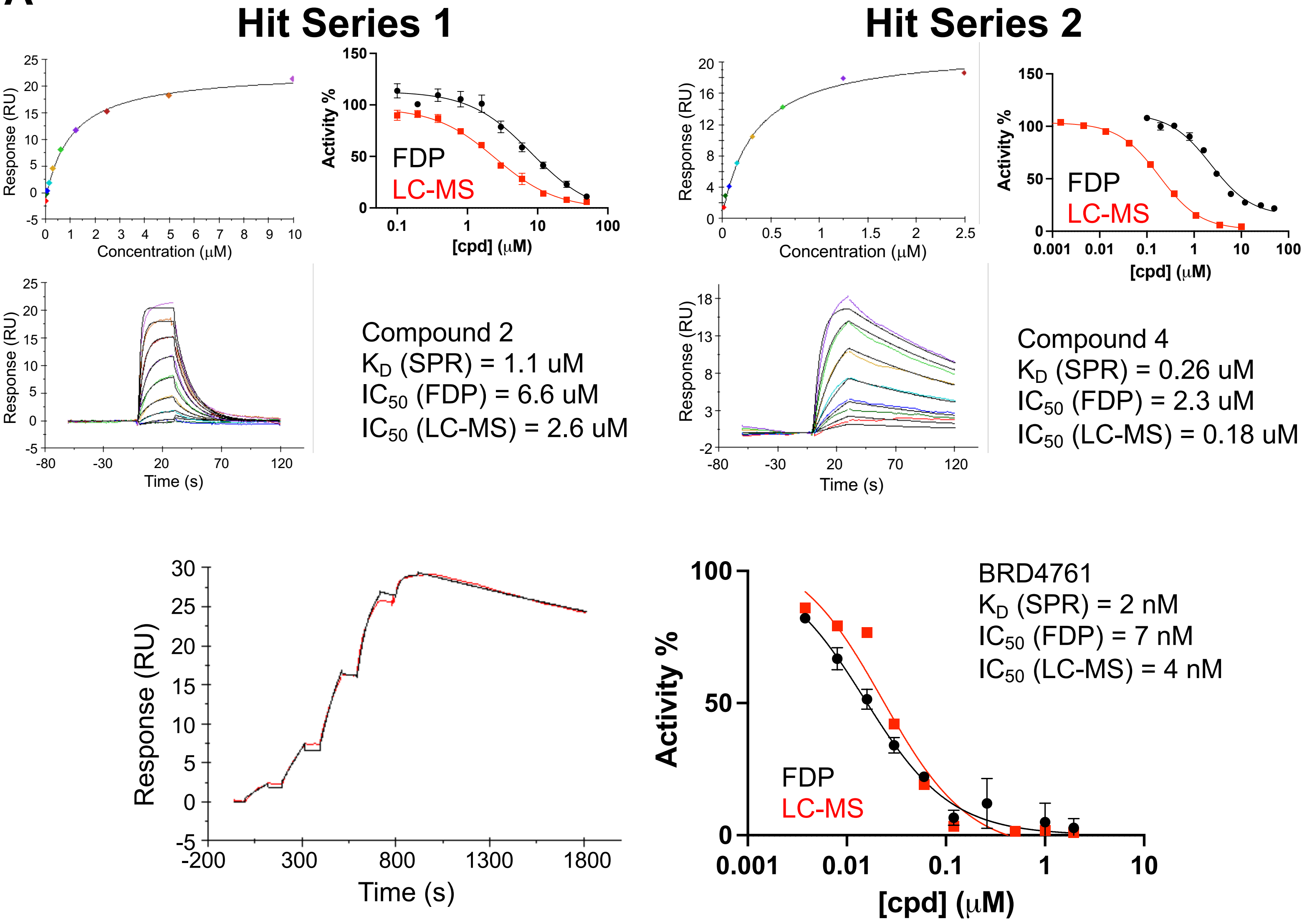


**B**

| Phosphatase[1] | Inhibition (10 uM) (%) | Phosphatase[1] | Inhibition (10 uM) (%) |
|---|---|---|---|
| PP1a[1] | 1 | PTPN11 (SHP-2)[1] | 5 |
| Lambda phosphatase[2] | -12 | PTPN22[1] | 22 |
| PP5[1] | -6 | PTPN4 (PTPMEG)[1] | -6 |
| ACP1 (LMPTP-A)[1] | 4 | PTPN6 (PTP1C, SHP-1)[1] | 10 |
| ACP1 (LMPTP-B)[1] | -1 | PTPN9 (MEG2)[1] | -4 |
| DUSP10 (MKP-5)[1] | -3 | PTPRB (PTPB)[1] | -2 |
| DUSP22[1] | -2 | PTPRC (CD45)[1] | 6 |
| DUSP3 (VHR)[1] | 1 | PTPRM[1] | -8 |
| HePTP[1] | -1 | TMDP[1] | -3 |
| PTPN1 (PTP1B)[1] | -9 | YohP[3] | 2 |

**Supplemental Figure 2. Lead Compound Characterization.** (A) Shown are the SPR data for lead compounds binding to PPM1D and enzymatic inhibition of PPM1D via the FDP (black) or LC-MS (red) assay. As outlined in **Figure 2**, the two hit series were combined to generate BRD4761. (B) Phosphatase Selectivity Panel Screening Results for BRD5049 run on the Eurofins Discovery PhosphotaseProfiler ([1]human; [2]lambda; [3]yersinia).

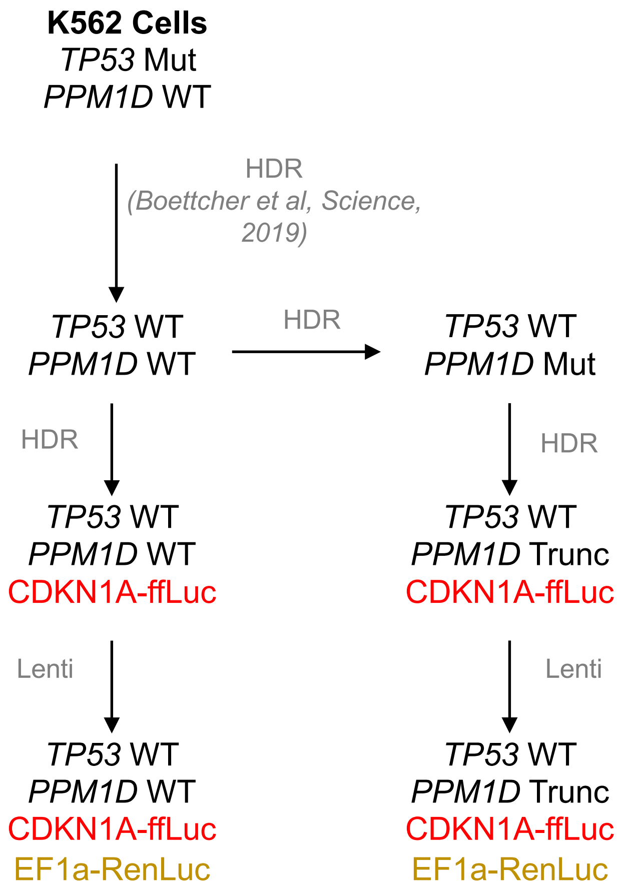


**Supplemental Figure 3. Schematic of Generation of p21-luciferase reporter system.**

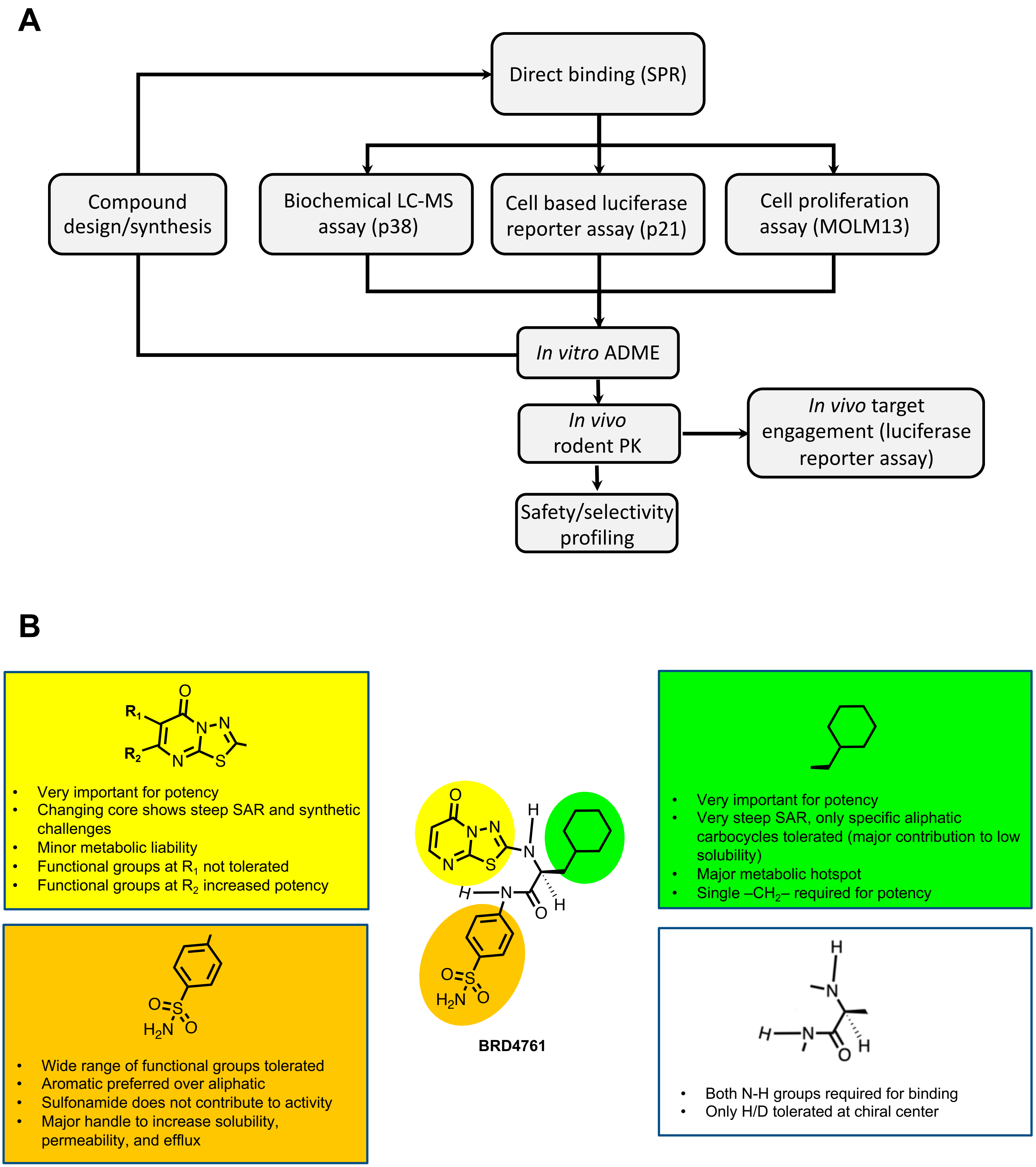


**Supplemental Figure 4. Lead Optimization.** (A) Overall experimental cascade for lead optimization. (B) Structure-Activity Relationships identified from lead compound optimization. Highlighted are how different parts of BRD4761 affect its activity.

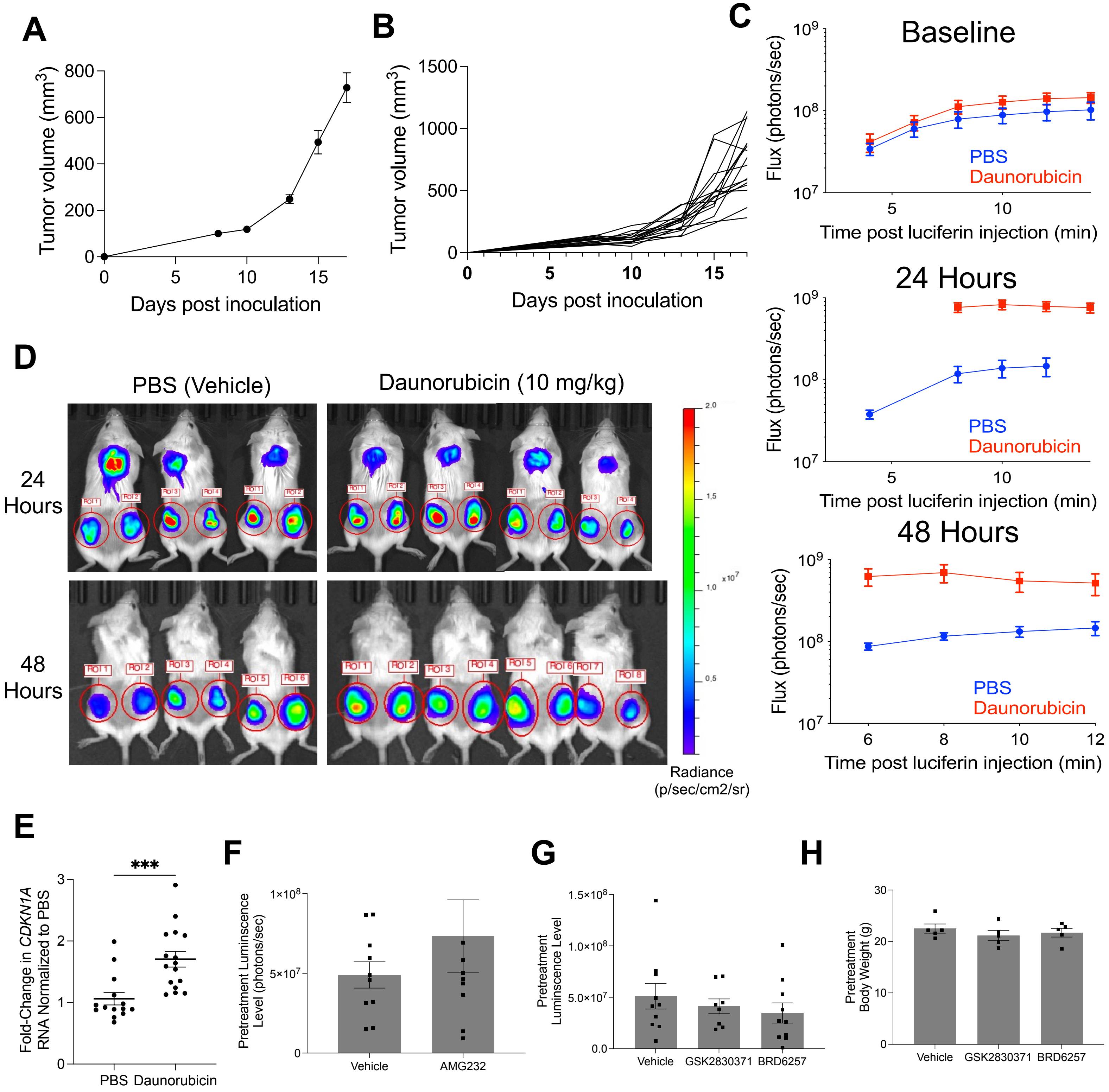


**Supplemental Figure 5. Pharmacokinetic optimization and pharmacodynamic evaluation *in vivo*.** Tumor growth after inoculation average across all recipients (A) and broken down by individual recipients (B). Luciferase flux after injection of luciferin reagents in mice treated with daunorubicin or phosphate buffered saline (PBS) control (C) and representative bioluminescent image of animals after treatment (D) (Note that the signal in the skull at 24 hours is a result of coelenterazine being oxidized by hemoglobin at the site of injection and is not due to a luciferase-based signal). (E) Quantitative PCR results of *CDKN1A* (p21) RNA levels in explanted tumors after treatment with PBS or daunorubicin. (F) Pretreatment tumor luminescence levels of animals treated with AMG232 or vehicle control. Pretreatment tumor luminescence levels (G) and bodyweight (H) of animals treated with vehicle, GSK2830371, or BRD6257.
